# NNMT drives innate sensitivity to NAMPT inhibition in YAP-dependent stem-like/ mesenchymal prostate cancer

**DOI:** 10.1101/2025.04.29.651153

**Authors:** Ágata Sofia Assunção Carreira, Marianna Ciuffreda, Nathakan Thongon, Charles M. Haughey, Adam Pickard, Sharon L. Eddie, Rebecca E. Steele, Elena Cerri, Romina Belli, Daniele Peroni, Elisa Facen, Irene Caffa, Moustafa Ghanem, Alessio Nencioni, Andrea Lunardi, Toma Tebaldi, Ian G. Mills, Alessandro Provenzani

## Abstract

Nicotinamide phosphoribosyltransferase (NAMPT) is the rate-limiting enzyme in the NAD+ salvage pathway and a promising therapeutic target in cancer. Resistance to NAMPT inhibitors, such as FK866, remains a key limitation to their clinical translation. While acquired resistance in cancer cell lines has been linked to target mutations, increased drug efflux and metabolic reprogramming, innate resistance mechanisms have been poorly studied. Addressing this gap is essential for identifying patient subgroups most likely to benefit from NAMPT-targeted therapies.

Advanced castration resistance prostate cancer (CRPC) lacks effective targeted treatments. Among its heterogeneous subtypes, stem cell-like CRPC (CRPC-SCL) is defined by androgen receptor (AR) independence, YAP/TAZ dependency, and mesenchymal traits. In this study, we identify the YAP/nicotinamide N-methyltransferase (NNMT) axis as a key mediator of innate sensitivity to FK866 in stem-like mesenchymal CRPC cells.

Using genetic and pharmacological models, we show that YAP or NNMT silencing rescues mesenchymal CRPC cells from FK866-induced apoptosis, endoplasmic reticulum stress, and NAD(H) loss. Metabolomic profiling confirmed that NNMT activity depletes nicotinamide, sensitizing cells to FK866. We further validated NNMT upregulation across clinical CRPC-SCL datasets, where it strongly correlates with mesenchymal and therapy-resistant phenotypes, as well as in murine prostate cancer cells with mesenchymal/stemness phenotypes.

In conclusion, we identify the YAP/NNMT axis as a determinant of innate sensitivity to NAMPT inhibition in prostate cancer. These findings support the use of NNMT as a predictive biomarker for NAD+-targeting therapies and provide mechanistic insight into a metabolic vulnerability of the CRPC-SCL subtype. Targeting the YAP/NNMT/NAMPT axis may represent a novel strategy for treating stem-like/mesenchymal, therapy-resistant prostate cancers.

**GRAPHICAL ABSTRACT:** 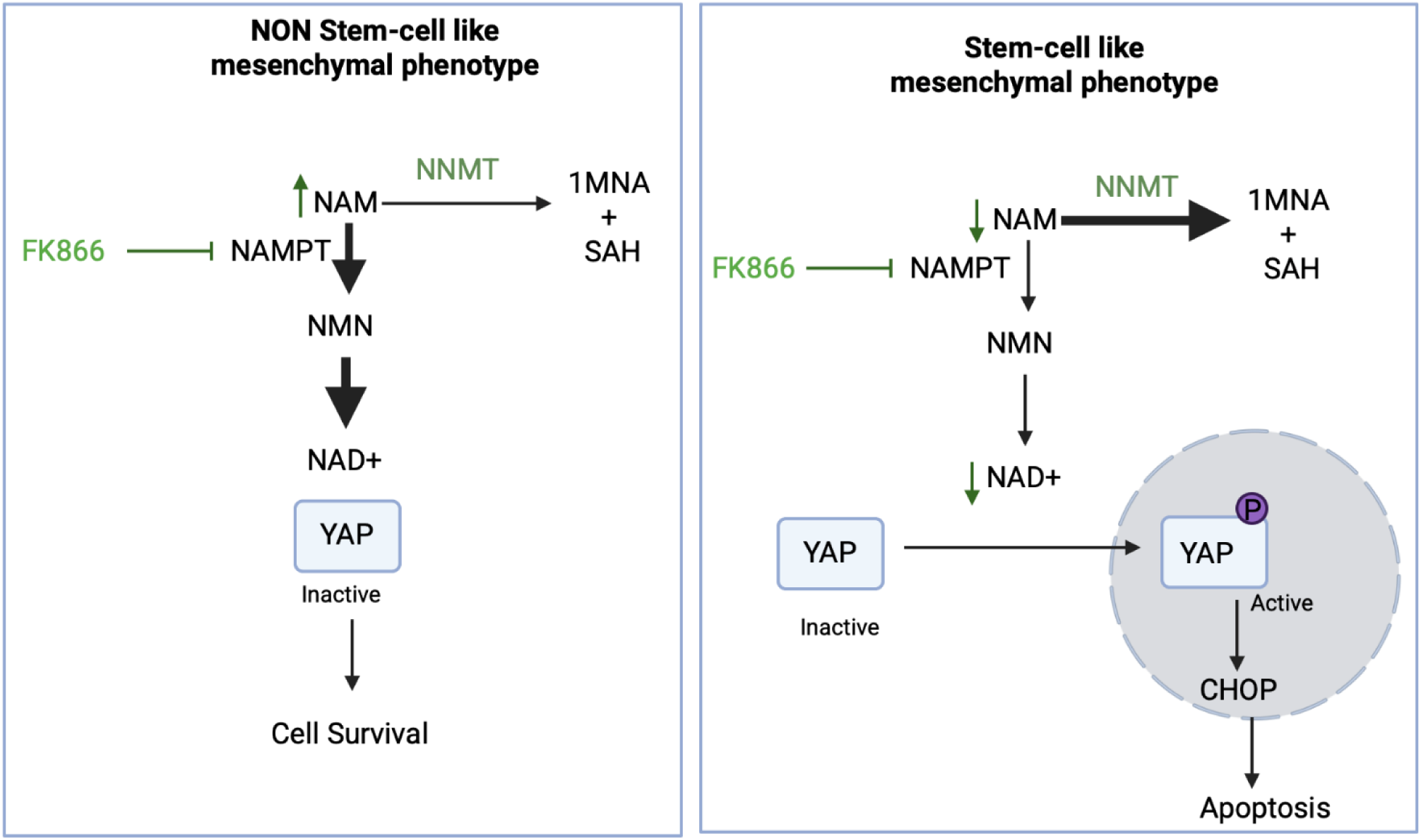

## INTRODUCTION

Malignant transformation encompasses a plethora of metabolic reprogramming processes to sustain cancer cells increased anabolic and catabolic needs^1^. The pyridine nucleotides nicotine adenine dinucleotide (NAD_+_) and its reduced form (NADH) play an essential role in cellular metabolism, signaling and DNA repair^2,3^.

Nicotinamide phosphoribosyl transferase (NAMPT) is the rate-limiting enzyme in the NAD^+^ biosynthesis, which has been found upregulated in several types of cancer cell lines and consequently emerged as a promising anti-cancer therapeutic target^4–7^, which supported extensive usage of NAMPT inhibitors, such as FK866 (APO866) in preclinical studies. Nonetheless, as often observed for small-molecules administration, acquired resistance to NAMPT inhibition in cancer cells has been reported. Resistance mechanisms to NAMPT inhibition can be sustained by target mutation, specifically in H191R and K342R^8^, as well as by increased ABC transporter-mediated drug efflux^9^, in human colorectal cancer cell lines, which were exposed for extended periods of time to FK866. We have also previously demonstrated that such long-term exposure to FK866 fully reprograms cell metabolism. Resistant triple-negative breast cancer cell lines have increased lactate dehydrogenase activity and mitochondrial function in order to counteract FK866-induced NAD(H) depletion^10,11^. Still, little is known about innate resistance mechanisms that would allow cancer cells to counteract NAMPT inhibitors’ toxicity without prior exposure. Such knowledge would contribute to the identification of patients and/or disease subgroups likely to benefit from NAMPT targeting either as single-use or combination therapy.

Prostate cancer (PC) is one of the most prevalent and frequently diagnosed cancers in men, with patients diagnosed with advanced high-risk PC often presenting as multiple-site metastasis^12^. Despite initial response to hormone deprivation therapy, a subset of patients will invariably develop resistance to the therapeutic regimen and develop castration-resistant PC (CRPC), for which targeted therapeutic options are not available^13^.

CRPCs often present with stem-cell like properties (CRPC-SCL) having undergone epithelial-to-mesenchymal transdifferentiation (EMT) potential or displaying mesenchymal phenotypes, which have been associated with tumor aggressiveness, chemo and radioresistance^14–16^. Recently, a pharmacogenomics *in-silico* study suggested FK866 efficacy in treating prostate tumors with stem cell-like properties^17^. The identification of mechanisms of resistance and/or sensitization to NAD^+^-depletion agents is crucial for the effective clinical application of NAMPT inhibitors, and can be particularly relevant for the treatment of stem-like CRPCs.

Here, we identified nicotinamide N-methyl transferase (NNMT) as a marker of a specific AP1/YAP-dependent subtype of CRPC with stem-cell-like properties (CRPC-SCL) and a sentinel player in the YAP-dependent molecular circuit promoting and sustaining the lethal progression of PCa. Mechanistically, we showed that YAP tumorigenic activity is linked to NNMT/NAMPT competition in mesenchymal stem-like CRPC cells. Importantly, by draining the intracellular pool of nicotinamide, NNMT activity determines the sensitivity of mesenchymal stem-like CRPC cells to NAMPT inhibition, which makes NNMT a promising predictive biomarker of the efficacy of NAD(H) depletion agents in CRPC-SCL patients. Our data was validated in publicly available prostate cancer patients’ datasets and in a preclinical model.

## MATERIAL AND METHODS

### Cell lines culture conditions

The PC cell lines PC3, LNCaP, VCaP and 22RV1 were obtained from the American Type Culture Collection (ATCC). PC3 cells silenced for YAP (PC3 shYAP) or NNMT (PC3 shNNMT), as well as the scramble PC3 cells (PC3 shSCR) were obtained as described below. PC3, LNCaP and 22RV1 cells were cultured in RPMI (Gibco), supplemented with 10% fetal bovine serum (FBS), 2 mM L-glutamine, and 100 U/mL penicillin-streptomycin (all from Lonza), while 22RV1 were cultured with DMEM (Gibco) containing the same supplementations. The supplemented RPMI culture media of PC3 shSCR, PC3 shYAP and PC3 shNNMT was additionally supplemented with 1 ug/mL of puromycin.

Murine prostate cancer cells (DVL3-PAR) were generated from tumors derived from the dorsal, ventral and lateral prostate lobes of a Pten−/−/trp53−/− Pb-Cre4 mouse^18^. DVL3-SCM were derived from the DVL3-PAR through culturing in Keratinocyte Serum Free Media (KSFM) + Bovine Pituitary Extract ((BPE) + Epidermal Growth Factor (EGF) (Thermo Fisher) + 2ng/ml Stem Cell Factor (SCF) (Stem Cell Technologies) +2ng/ml Leukaemia Inhibitory Factor (LIF) (Sigma) + 1ng/ml granulocyte-macrophage colony–stimulating factor (GM-CSF) (Miltenyi Biotec)+ 2mM L-glutamine (Gibco). Cells were kept under humidified conditions with 5% CO2 at 37 °C.

### Drug treatment and in vitro cell viability

Cell lines were treated with FK866 (sc-205325, Santa Cruz Biotechnology). For *in vitro* drug sensitivity assessment, cells were treated for 48 hours with FK866 and the OzBlue Cell Viability kit (OzBiosciences) was used to measure the compound toxicity after 2 hours of incubation, according to the manufacturer’s instructions.

### Quantification of NAD(H) and ATP levels

NAD(H) and ATP levels were quantified after 48 hours of FK866 exposure. Intracellular NAD(H) was measured with the NAD/NADH-Glo (G9071, Promega), while ATP levels were measured with the CellTiter-Glo Luminescent Cell Viability Assay (G7571, Promega), according to the manufacturer’s protocol. Luminescence data were normalized to protein content in the cell lysates under the same conditions, using the Bradford assay (Bradford Reagent, Sigma).

### Apoptosis (Caspase 3/7 activation) assay

Activation of apoptosis was assessed using the CellEventTM Caspase 3/7 Green (Life Technologies, C10432), which generates bright green fluorescence upon caspase 3/7 activation. LNCaP, PC3, PC3 shSCR and PC3 shNNMT were seeded into tissue cultured transparent clear-bottom 96-well plates and allowed to adhere for 24 hours. Cells were then cotreated with FK866 at the indicated concentrations and with the CellEventTM Caspase 3/7 Green reagent. After 72 hours, both cell number (brightfield channel) and green fluorescent (FITC channel) signals were detected with IncuCyte® S3 Live-Cell Analysis System (Sartorius). Images were analyzed for the number of green objects (apoptotic cells) per well. Results are presented as a fold change in the respective vehicle (DMSO) conditions.

### RNA extraction and Real-Time quantitative PCR (RT-qPCR)

Total RNA extraction was accomplished by aqueous-organic phase separation with TriZOL reagent (Invitrogen). cDNA synthesis (RevertAid RT Reverse Transcription Kit, K1691, ThermoScientific) was performed on the extracted RNA, according to the manufacturer’s instructions. RT-qPCR runs were performed in technical triplicates using the ExcelTaq™ 2X qPCR Master Mix (SMOBIO) on a CFX96 Real-Time PCR Detection System (Bio-Rad). The sequences of the primers used are described in **Supplementary Material 1**. ΔCq method was used for amplified mRNA quantification with RPLP0 as the housekeeping gene.

### Protein extraction and western blot analysis

RIPA lysis buffer supplemented with Protease Inhibitor Cocktail (ThermoScientific) was used to lyse the collected samples. After clarification, protein extracts were quantified with the Pierce™ BCA Protein Assay (23225). Equal amounts of proteins (20 µg) were loaded in SDS-polyacrylamide gel, followed by the transference to a PVDF membrane. Primary and secondary antibodies used are listed in **Supplementary Material 1**. The chemoluminescent signal was detected with Amersham ECL Select from GE Healthcare at ChemiDoc (BioRad). A representative western blot of three independent biological experiments is shown.

#### Lentiviral particles (LVPs) production and infection

Lentiviral vector particles (LVP) for shYAP and shNNMT were produced in HEK293T cells, using pLKO vector as the transfer plasmid. HEK293T cells were transfected with both a Δ891 and a VSV-G encoding vector along with the shSCR/shYAP/shNNMT plasmids. 72 hours post-transfection, the viral supernatants were collected and filtrated. Following the quantification with the PERT assay, 1 Reverse Transcription Unit (RTU) was used to infect PC3 cells. Selection with puromycin (1 µg/mL) was performed on PC3 cells stably transduced with shSCR/shYAP/shNNMT.

### Soft agarose (colony formation assay)

Colony formation in anchorage-independent conditions was evaluated with the soft agarose assay. The assay was carried out in 6-well plates with a double coating of agarose: the base agarose layer presented 0,7% of agarose, while the upper coat containing the cells presented 0,35% of agarose. For the top layer, 5000 cells per condition were seeded. Cells were covered with 1mL of complete media and treated with different concentrations of FK866 every 3 days from day 1, for 14-15 days. Total colony number was quantified with Operetta and Harmony 3.5.2 software.

### Proteomics of PC3 shYAP

PC3 shSCR and shYAP were seeded and treated with FK866 or vehicle conditions for 48 hours, after which they were collected and processed for proteomics analysis.

### Sample preparation

Digested samples were separated using an Easy-nLC 1200 system (Thermo Scientific). A 30 cm reversed-phase C18 column (inner diameter 75 µm, 1.7µm particle size, MSWIL, the Netherlands) heated at 40 °C was employed for peptide separation, with a two-component mobile phase system consisting of 0.1% formic acid in water (buffer A) and 0.1% formic acid in 80% acetonitrile (buffer B). Peptides were eluted using a gradient of 5% to 25% over 55 minutes, followed by 25% to 40% over 10 minutes, by 40% to 98% over 10 minutes, and finally kept at 98% over 10 minutes. The flow rate was set to 200 nL/min. Samples were injected into an Orbitrap Fusion Tribrid mass spectrometer (Thermo Scientific, San Jose, CA, USA), and data were acquired in data-dependent mode (2100 V). The temperature of the ion transfer tube was maintained at 200 °C. Full scans were performed at a resolving power of 120,000 FWHM (at 200 m/z) with an AGC target of 1 × 10⁶. The mass range of 350–1100 m/z was surveyed for precursors, with the first mass set at 140 m/z for fragments. Each full scan was followed by a set of MS/MS scans (HCD, collision energy of 30%) with a 3-second cycle time and a maximum injection time of 150 ms in the ion trap, along with an AGC target of 5 × 10³. A dynamic exclusion filter was applied every 40 seconds with a ±5 ppm mass tolerance. Data were acquired using Thermo software Xcalibur (version 4.3) and Tune (version 3.3). During the project, QCloud^19^ was used to monitor the instrument’s longitudinal performance. Peptide searches were conducted using Proteome Discoverer 2.2 software (Thermo Fisher Scientific) against the Homo sapiens FASTA file (UniProt, downloaded in March 2023) and a database containing common contaminants. Proteins were identified using the MASCOT search engine with a mass tolerance of 10 ppm for precursors and 0.6 Da for products. Trypsin was selected as the enzyme, allowing a maximum of 5 missed cleavages. Static modification of carbamidomethyl (C) and variable modifications of oxidation (M) and acetylation (protein N-term) were included in the search. False discovery rate filtering was applied, with a threshold set to <0.05 at the PSM, peptide, and protein levels. Results were filtered to exclude potential contaminants and proteins with fewer than two peptides.

### Computational analysis

MS downstream analysis was conducted using the ProTN proteomics pipeline available at www.github.com/TebaldiLab/ProTN and www.rdds.it/ProTN. In summary, peptide intensities were log_2_-transformed, normalized using median normalization, and summarized into proteins through functions in the DEqMS Bioconductor package^20^. Imputation of missing intensities was performed using the PhosR package^21^. Differential analysis was carried out with the DEqMS package, where proteins with an absolute log_2_ fold change (FC) greater than 0.75 and a p-value less than 0.05 were deemed significant. Functional enrichment analysis of differentially expressed proteins was conducted using the ClusterProfiler function EnrichGO^22^. Enriched terms with a p-value lower than 0.05 were considered significant. The mass spectrometry proteomics data have been deposited to the ProteomeXchange Consortium via the PRIDE partner repository with the dataset identifier PXD056852 and 10.6019/PXD056852^23^ (**Supplementary Material 2**).

### RNA expression data analysis

RNA sequencing (RNA-seq) data for cell lines and organoids were obtained from ^24^, while patients and patient-derived xenograft (PDX) data were accessed through Prostate Cancer MDA PC PDX dataset from the cBioPortal. For the analysis of NNMT expression across cell lines, organoids, and our mouse models, regularized logarithm (rlog) normalized read counts generated using DESeq2 were utilized. In contrast, for patients and PDX samples, RSEM batch-normalized counts were employed. Statistical significance of NNMT expression in our mouse models was assessed using a two-tailed t-test, while the Wilcoxon rank-sum test was employed to determine stastical significance for cell lines, organoids, patients and PDX models. To integrate the expression data from cell lines, organoids, and PDX models, the COMBAT algorithm from the sva R package v3.52.0 was applied to correct batch effects. Dimensionality reduction and visualization of the integrated data were conducted using Uniform Manifold Approximation and Projection (UMAP). The epithelial-to-mesenchymal transition (EMT) score was calculated following the 76 gene signature (76GS) method described in ^25^.

### Xenografts

C57BL/6 male mice (8 weeks old) were obtained from Harlan, UK and allowed to acclimatize to their environment for a minimum of one week prior to experiment. Mice were anaesthetized using isoflourene prior to engraftment, with treatment maintained throughout the procedure. Using a 23G needle, mice were inoculated subcutaneously with 2 x 10^6^ DVL3-PAR or DVL3-SCM cells which had been suspended in 100μl of sterile PBS and Matrigel (Corning) at ratio of 1:1. After inoculation mice were carefully monitored until they regained full consciousness. Mice weights and tumor volumes were measured twice a week until the predetermined endpoint of the experiment was reached.

Tumor bearing mice were sacrificed at the indicated timepoints, tumors excised and divided. A portion of the tissue was snap frozen for protein, snap frozen for RNA, stored in RNA later (Sigma) for 24hrs before long term storage at-80 °C and a final portion was fixed in 4% buffered formalin (Sigma Aldrich, UK). Tumors collected in Manchester were fixed in 4% buffered formalin (Sigma Aldrich, UK) for 24 hours or collected in media for tumor disaggregation. The formalin fixed tumor samples were transferred to 70% ethanol before being processed into FFPE blocks, cut using a microtome at a thickness of 3μm and mounted onto glass slides by the precision medicine center.

### RNA extraction for RNA seq analysis

RNA from FFPE mouse tumor tissue was extracted by colleagues in Prof Tim Illidge’s laboratory at Manchester University. Tissue sections, 2-4 (10μm) sections were transferred into RNAase free microcentrifuge tube. The paraffin was removed by incubating in xylene and ethanol. For lysis and total RNA purification, digestion buffer and proteinase-K were added to the samples as per the manufacturer’s instruction (Norgen kit). The samples were spun briefly followed by transferring the supernatant to a new RNAase free microcentrifuge tube. The RNA-containing tubes were incubated for 15 minutes at 80oC. The lysates were then passed through an RNA purification microcolumn and centrifuged for 1min at 14,000rpm. The microcolumns were washed according to the manufacturer’s instruction and the RNA was eluted using the Elution solution (Norgen Kit).

### RNA library preparation, sequencing, and analysis

Libraries were generated from an input RNA concentration of 500ng per sample using the using the Quant-Seq 3’mRNA SEQ Library Prep Kit (Lexogen). The adapted version of this protocol was used for RNA samples which had been extracted from FFPE samples as per manufacturer’s instructions. Pooling and sequencing of the libraries on the NextSeq was performed in the CCRCB genomic core facility and reads were aligned to the MM10 genomic reference. Mouse orthologs were converted to their human equivalent using the online tool accessed at https://biit.cs.ut.ee/gprofiler/gorth.ci. Differential gene expression analysis was performed using the open source, web-based platform Galaxy which determines the differentially expressed genes using the DESeq2 tool.

### Soft agar colony formation assay

Anchorage independent growth was performed as previously described^26^. Shortly, DVL3-PAR/SCM cells were seeded in a 24 well plate in 0.35% agarose/DMEM solution, and kept for 3 weeks with media change every 4 days. A representative image is shown.

### Analysis of metabolites

#### Sample preparation

LNCaP, PC3 shSCR and PC3 shNNMT cells were treated for 48 hours with 10 µM concentration of FK866. After removal of the culture medium, cells were washed with 1 mL of cold 150 mM ammonium acetate. The extraction solvent (80 % pre-chilled methanol) was added directly to each well, and the plates were left for 20 min at −80 °C. The cells were scraped, harvested in 1.5 ml Eppendorf tubes, and vortexed for 2 min. The solution was then centrifuged at 14,000 g for 10 min at 4 °C to remove the protein pellet. The supernatants were transferred to a new tube, dried in a SpeedVac vacuum and stored at −80 °C. Before the analysis, the dried metabolite samples were reconstituted in 50 μL of 3% acetonitrile/0.1% formic acid aqueous solution and thoroughly mixed. The samples were centrifuged for 2 minutes at 14,000 g, and the supernatants were subjected to LC/MS analysis. Seven independent biological replicates were prepared.

To prepare the standards for the calibration curves of NAM and NAD+, seven serial drug dilutions, ranging from 6.4nM to 100μM with 3 technical replicates each, were resuspended in 3% acetonitrile/0.1% formic acid aqueous solution starting from a 10 mM stock solution. The lower limit of detection (LOD) was determined by comparing the signals from a set of samples with known low concentrations of analyte with those of control samples (DMSO). The minimum signal-to-noise <u>ratio</u> for the LOD was set to the ratio of 3:1.

#### LCMS Data acquisition

An amount of 10 µl of each standard and sample was injected into an Orbitrap Fusion™ Tribrid™ mass spectrometer (Thermo Scientific) coupled with a Dionex Ultimate 3000RS chromatography system (Thermo Scientific). A Kinetex® 2.6 µm Polar C18 column (Phenomenex, 150 × 2.1 mm) was used for separation. The column temperature was set at 40 °C. The LC method consisted of a gradient from 1 to 15%B (B: acetonitrile with 0.1% formic acid; A: water with 0.1% formic acid) over 5 min, from 15 to 100%B over 2 min, followed by 3 min at 100% at a flow rate of 0.2 ml/min. The MS spray voltage was set at 3500 V with an ion transfer tube temperature of 300 °C. Sheath and auxiliary gasses were set at 20 and 5 Arb, respectively. The MS data were acquired in full scan mode at 120,000 FWHM (200 m/z), in the scan range of m/z 100–1000, with maximum injection time of 50ms and the AGC set to 4e5. Standards were analyzed in triplicates.

## Data analysis

The area under the peak of each precursor ion and the total ion current (TIC) were extracted using Skyline (MacCoss Lab Software) for standards and samples^27^. The software derived precise *m*/*z* values as well as isotope distributions for each precursor. Each analyte was also investigated for common adducts, [M+H]+, [M+K]+, [M+NH4]+, and [M+Na]+ ions. The normalization of the area under the peak of each ion by the TIC of the sample was performed to account for variations in sampling volumes across conditions. To build the calibration curve, Skyline calculates the mean of the observed normalized peak areas of the standards at each concentration level. The log-transformed value of the total area of precursor ions of each concentration level was then plotted as a function of the concentration to build the standard curves for each compound. Finally, standard linear regression was performed to obtain the linear predictors that were used to compute the concentration value of the compound.

## Statistical analysis

The presented data was expressed as mean ± standard error of mean (SEM) of three independent biological experiments conducted in technical triplicates, unless indicated otherwise. Statistical significance between control and test groups was determined using individual Student’s t-test, considering values of *p < 0.05, **p < 0.01, and ***p < 0.001 as significant. Data was plotted by GraphPad Prism 8 software.

## RESULTS

### PC cell line PC3 shows high sensitivity to FK866

To characterize innate resistance mechanisms to NAMPT inhibition in cancer cells, we assessed cellular toxicity to the NAMPT inhibitor FK866 in a panel of cancer cell lines. We observed that the PC cell lines PC3 and LNCaP presented different sensitivity to FK866. PC3 cells were found to be more sensitive to FK866 than LNCaP, showing reduced cell viability at doses ineffective in LNCaP cells (**Figure 1A**). Exposure to FK866 for 48 hours decreased both NAD(H) and ATP levels in PC3, in a dose-dependent manner (**Figure 1B-C**), while LNCaP maintained stable NAD(H), even when exposed to higher concentrations of the small molecule, and ATP levels (**Figure 1B-C**). To assess the impact of NAD(H) depletion in apoptosis induction, PC3 and LNCaP cells were exposed to FK866 for 72 hours, in the presence of fluorogenic substrates for activated Caspase 3/7^24^. Incucyte® analysis revealed an increased apoptosis activation in both PC3 and LNCaP cells treated with the NAMPT inhibitor, which was more efficient in PC3 cells (**Figure 1D**). A broader analysis of FK866 efficacy in a PC cell lines panel demonstrated a strong negative correlation (Pearson r = - 0,9807; p-value = 0.0032) between NAMPT expression and the viability after FK866 exposure, suggesting that higher expression/dependency on NAMPT increases PC cells sensitivity to NAMPT inhibition (**Figure 1E**). Indeed, PC cells with stem-like properties and differentiation potential (epithelial to mesenchymal transdifferentiation) were previously described to present increased sensitivity to FK866^17^. Accordingly, PC3 cells, that show the most mesenchymal phenotype, are characterized by a very low expression of *CDH1* (E-Cadherin) and high expression of *CDH2* (N*-Cadherin), VIM* (*Vimentin*), *SNAIL1*, and *ZEB1* (**Figure 1F**). Differently, LNCaP, VCaP and 22RV1 cell lines showed high levels of *CDH1* and low levels of the mesenchymal markers compared with PC3 cells, in line with their marked epithelial phenotype (**Figure 1F**).

**Figure 1.**
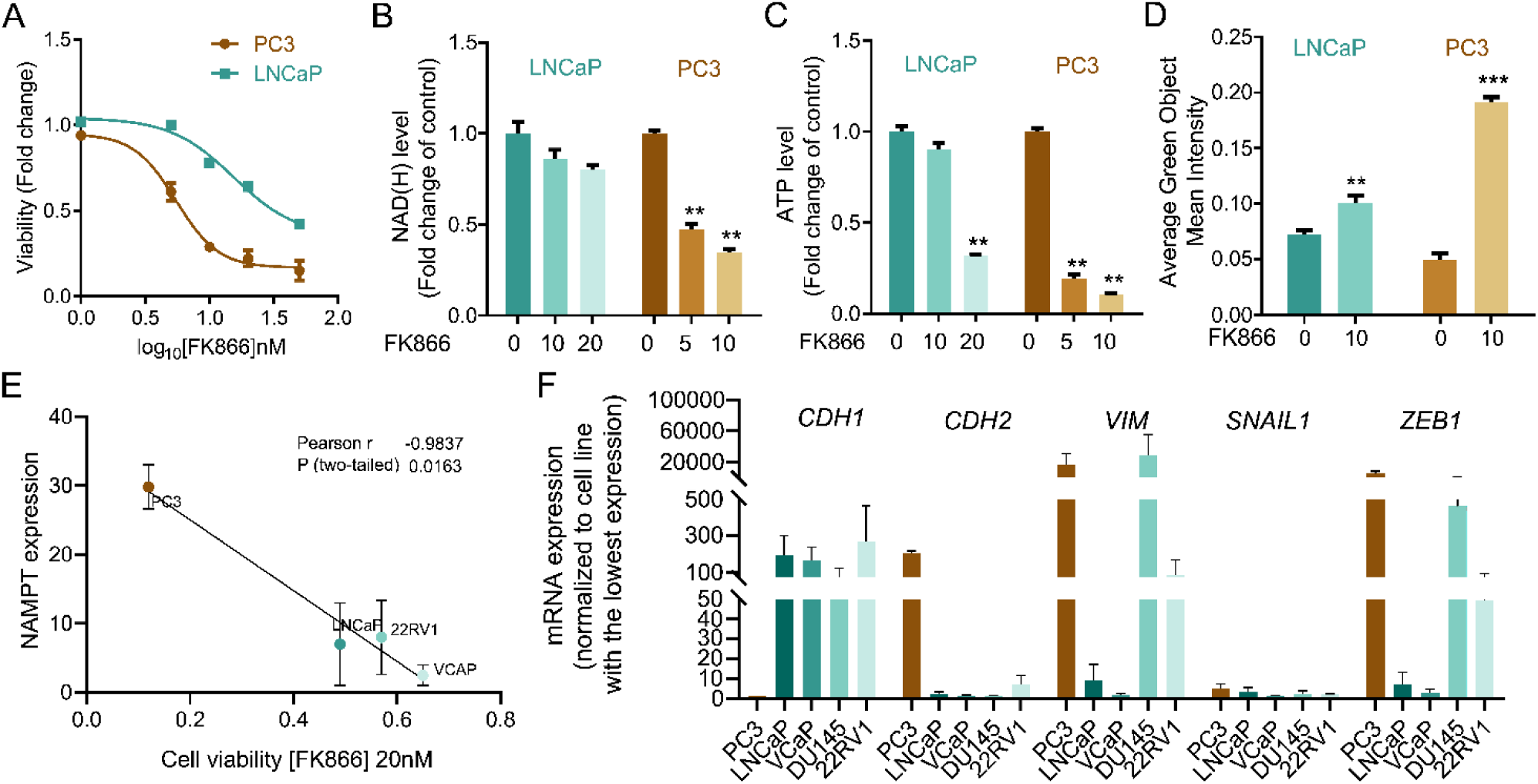
: PC3 cells are more sensitive to NAMPT inhibition than LNCaP. **(A)** Viability curve of PC3 and LNCaP cells after exposure to FK866 for 48 hours. Viability calculated as a fold change of the DMSO condition. **(B)** NAD(H) and **(C)** ATP levels in PC3 and LNCaP cells upon exposure to FK866 at concentrations of 5, 10 or 20 nM for 48 hours. Data presented as a fold change of the control condition for each cell line. **(D)** Quantification of caspase 3/7 activation in PC3 and LNCaP upon 72 hours of FK866 exposure at 10 nM. Data were obtained with Incucyte, and results are presented as a fold change to non-treated condition for each cell line. **(E)** Linear regression of NAMPT transcript expression in function of the cell viability of prostate cancer cell lines exposed to FK866 at 20 nM. The Pearson correlation coefficient was used to measure the linear correlation between the two data sets, using three independent biological replicates. **(F)** RT-qPCR analysis of epithelial (*CDH1*) and mesenchymal genes (*CDH2*, *VIMENTIN* - *VIM*, *SNAIL1* and *ZEB1*) in a panel of prostate cancer cell lines. The mRNA expression, calculated with the ΔCt method, was normalized to the cell line with the lowest expression (highest Ct value) of that gene.For all the experiments, three independent biological replicates were performed and used for statistical analysis. Repeated measures one-way ANOVA were used to calculate statistical significance (ns-not significant, *P < 0.05, **P < 0.01, ***P < 0.001) between DMSO (FK866 0nM) and experimental conditions.

Overall, these data indicate a correlation between NAMPT expression and EMT phenotypes, and sensitivity to NAMPT inhibition in prostate cancer cell lines. Still, further characterization of the cellular models is needed to unravel mechanisms of innate resistance to FK866.

### YAP is a mediator of FK866 sensitivity in PC3 cells

The classification and molecular characterization of CRPC subtypes is a major challenge in the prostate cancer field. Recently, a classification of PC cell lines, organoids, and patient-derived xenografts (PDXs) based on their chromatin profiles and transcription factors’ dependencies has been proposed, highlighting four CRPC subtypes sustained by AR (“CRPC-AR”) or Wnt pathway dependency (“CRPC-Wnt”), neuroendocrine phenotype (“CRPC-NE”), and an AR-independent stem-cell like phenotype (“CRPC-SCLC”)^24^. While the LNCaP cell line is classified as “CRPC-AR”, PC3 cells are “CRPC-SCLC”. This subtype is driven by activated protein-1 (AP-1) transcription factors and further sustained by interactions with the YAP/TAZ proteins^24^. Thus, we decided to evaluate YAP response to different levels of NAD(H), by assessing its phosphorylation status by western blotting in PC3 and LNCaP treated with FK866. Phosphorylated YAP has a cytoplasmic localization and is inactive, while dephosphorylation drives YAP nuclear translocation and initiation of YAP-dependent transcriptional program. Differently from LNCaP, FK866 exposure decreased YAP phosphorylation at serine 127 in PC3 cells, without modulation of total YAP levels (**Figure 2A**). Consistently, the *bona fide* YAP target genes *Connective tissue growth factor* (*CTGF/CCN2)* and *Cysteine-rich angiogenic inducer 61* (*CYR61/CCN1)* were dose-dependently upregulated in PC3 cells treated with FK866, whereas there was no impact on LNCaP (**Figure 2B-C**).

**Figure 2.**
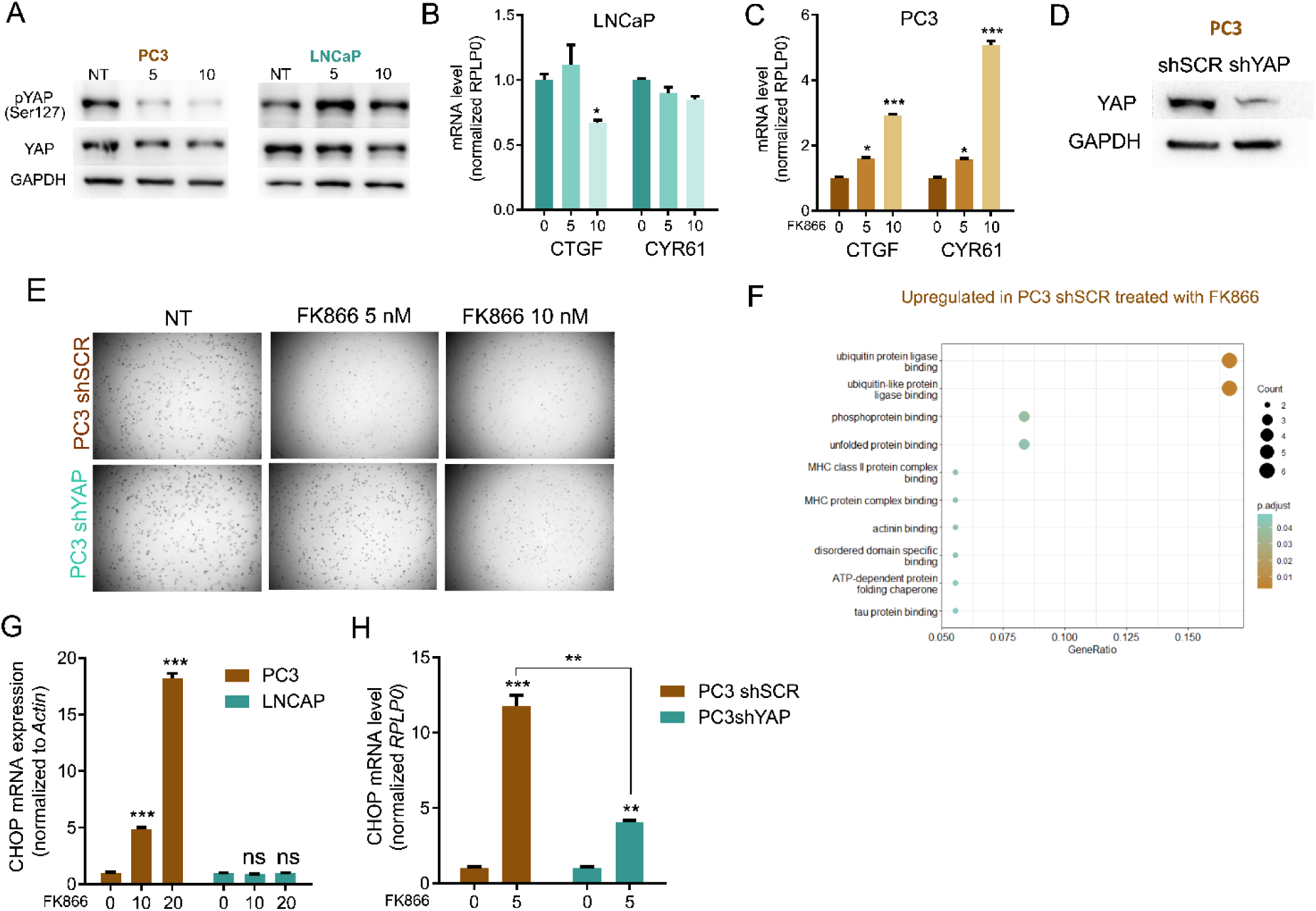
NAD(H) depletion activates YAP and CHOP in PC3, but not LNCaP cells. **(A)** Western Blot analysis for the expression of phosphorylated YAP at Serine 127 (p-YAP Ser127) and total YAP protein in PC3 and LNCaP cells, after the treatment for 48 hours with FK866 at concentrations 5 or 10 nM. Actin was used as housekeeping. **(B)** and **(C)** mRNA levels of YAP targets *CTGF* and *CYR61* in PC3 and LNCaP cell lines, respectively, after the exposure for 48 hours with 5 or 10 nM concentrations of FK866. RPLP0 was used as the housekeeping gene. Three independent biological replicates were performed. **(D)** Western Blot for the expression of YAP protein in PC3 cells stably transfected with the scramble (shSCR) or the short-hairpin YAP silencing construct (shYAP). GAPDH was used as housekeeping. (E) Soft agarose assay of PC3 shSCR and shYAP cells treated with 5 or 10 nM of FK866, every 2-3 days for 2 weeks. Images acquired with Operetta and Harmony 3.5.2 software. **(F)** Gene set enrichment analysis (GSEA) of proteins upregulated in PC3 shSCR cells treated with FK866 (5nM) for 48 hours. All the significant upregulated proteins (p-value above 0,05) compared with the DMSO treated conditions were considered. There independent biological replicates were considered for statistical analysis. **(G)** mRNA levels of CHOP in PC3 and LNCaP cell lines after exposure for 48 hours with 5, 10 or 20 nM concentrations of FK866, **(H)** mRNA levels of CHOP in PC3 shSCR and PC3 shYAP cell lines, treated with 5nM concentration of FK866 for 48 hours. RPLP0 was used as the housekeeping gene. Three independent biological replicates were performed. Repeated measures one-way ANOVA was used to calculate statistical significance (ns-not significant, *P < 0.05, **P < 0.01, ***P < 0.001) between DMSO (FK866 0nM) and experimental conditions.

The stable silencing of YAP in PC3 cells (PC3 shYAP), confirmed by western blotting (**Figure 2D**), reduced their sensitivity to FK866 (**Figure 2E**) compared with the one of PC3 cells transfected with a scrambled short-hairpin RNA (PC3 shSCR), which suggests that YAP is a mediator of sensitivity to FK866. To gain deeper insights into the molecular pathway inducing cell death of PC3 cells, activated by YAP upon NAD(H) depletion, we performed label-free quantitative proteomics on PC3 cells treated with 5 nM FK866 for 48 hours. These cells presented an upregulation of proteins related to “ubiquitin(-like) protein ligase binding” and “unfolded protein binding”, which suggested induction of the protein quality control system (**Figure 2F**). Indeed, C/EBP-homologous protein (CHOP), was found upregulated in PC3 cells treated with FK866 but not in LNCaP cells (**Figure 2G**). As expected, CHOP expression was also induced in PC3 shSCR exposed to FK866, and partially rescued by YAP silencing (**Figure 2H**). Of note, PC3 cells treated with FK866 showed downregulation of proteins associated with “mitochondrial matrix”, “mitochondrial translation”, “mitochondrial gene expression” and “mitochondrial ribosome” (**Supplementary Figure 1A**), supporting the importance of increased mitochondrial mass and function to counteract FK866 toxicity, as previously reported^2^. Overall, these data indicate that YAP regulates the CHOP-dependent stress response in PC3 cells arising from NAD(H) depletion.

### NNMT is downregulated in PC3 shYAP

To gain further molecular insights into the role of YAP as a mediator of FK866 sensitivity, we performed label-free quantitative proteomics on PC3 cells stably silenced for YAP (PC3 shYAP). Proteomics data showed YAP downregulation in PC3 shYAP (log_2_ of fold change (FC) =-1.89; p-value =3,5 x 10^-5^), confirming the efficacy of the silencing (**Figure 3A**). We identified 3853 proteins within this dataset, of which 329 were differentially expressed in PC3 shYAP compared with PC3 shSCR (p-value lower than 0.05 – **Supplementary Material 1**). Both up-and down-regulated proteins in PC3 shYAP were independently subjected to gene enrichment analysis (GEA). Unsurprisingly, the 172 down-regulated proteins in PC3 shYAP exposed terms associated with YAP-regulated cell mechanics’ processes, such as “actomyosin structure organization”, “cell substrate/cell-matrix adhesion” and “regulation of actin filament organization” (**Supplementary Figure 1B**).

**Figure 3.**
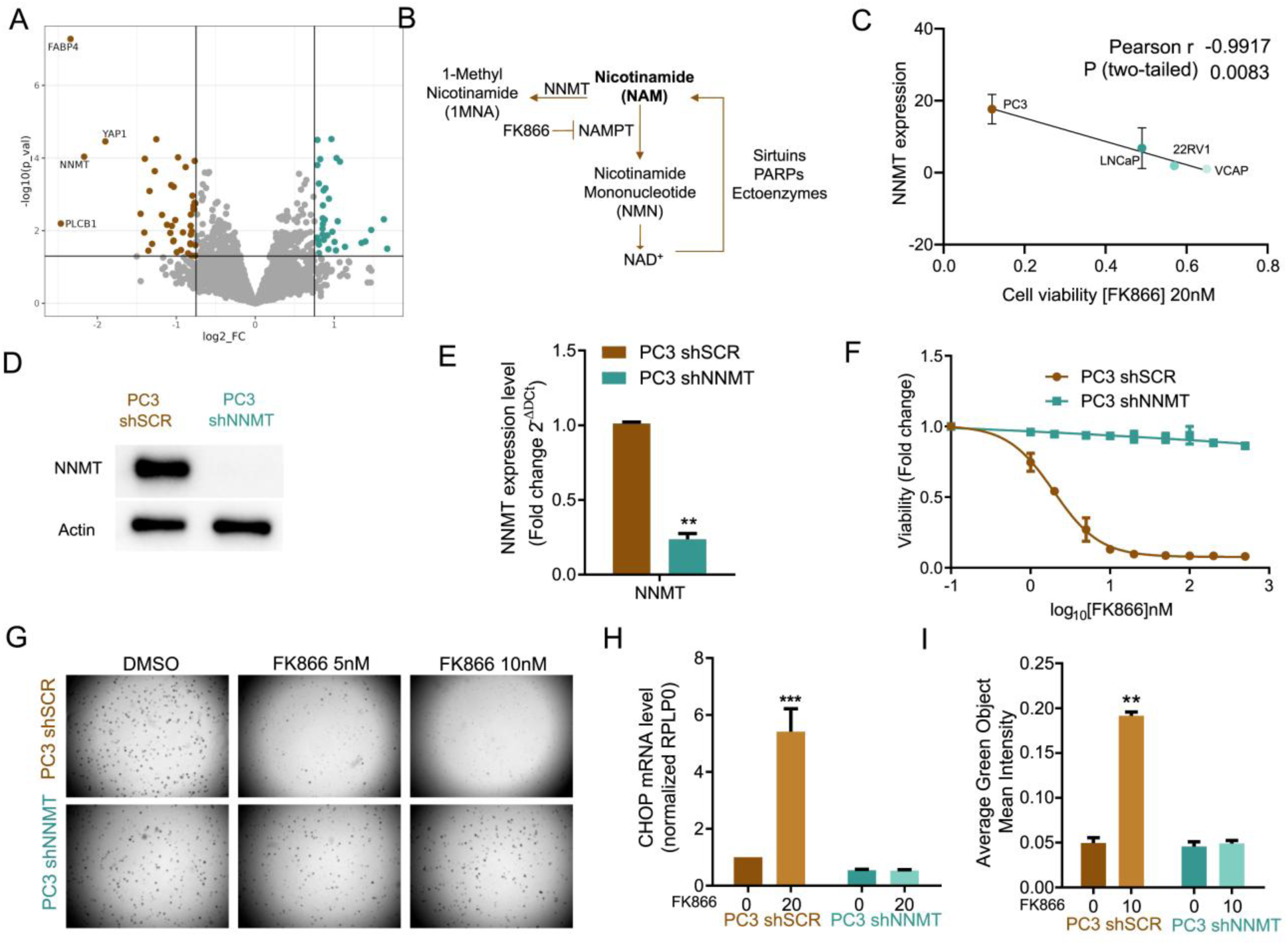
NNMT is a mediator of FK866 sensitivity in PC3 cells in vitro. (**A**)Volcano plot of log_2_ of the fold change (FC) of the protein expression, versus-log_10_ of the p-value (p-val) calculated for four independent biological replicates. The threshold was defined as-log_10_(p-val) above 1.3, with log_2_FC respectively above 1.7 of fold change for upregulated proteins, and 0.6 of fold change for downregulated proteins. Dots colored in brown and in blue represent respectively the down and up-regulated proteins in PC3 shYAP **(B)** Schematic representation of nicotinamide (NAM) synthesis and consumption pathways. **(C)** Linear regression of NNMT transcript expression in function of the cell viability of prostate cancer cell lines exposed to FK866 at 20 nM. The Pearson correlation coefficient was used to measure the linear correlation between the two data sets, using three independent biological replicates. **(D)** Western Blot and **(E)** RTqPCR analysis for the expression of NNMT in PC3 cells stably transfected with the scramble (shSCR) or the short-hairpin NNMT silencing construct (shNNMT). Actin was used as housekeeping. **(F)** Viability curve of PC3 shSCR and PC3 shNNMT cells after exposure to FK866 for 48 hours. Viability calculated as fold change of the DMSO condition. Three independent biological replicates were performed. **(G)** Soft agarose assay with PC3 shSCr and PC3 shNNMT exposed to FK866 at 5 and 10 nM (or DSO control condition) for about 2 weeks. Media supplemented with the small molecule substituted every 2-3 days. Representative images acquired with Operetta and Harmony 3.5.2 software. **(H)** Quantification of caspase 3/7 activation in PC3 shSCr and PC3 shNNMT upon 72 hours of FK866 exposure at 10 nM. Data were obtained with Incucyte, and results are presented as a fold change to non treated condition for each cell line. **(I)** mRNA levels of CHOP in PC3 shSCR and PC3 shNNMT cell lines after exposure for 48 hours with 20 nM of FK866. RPLP0 was used as housekeeping gene. Repeated measures one-way ANOVA were used to calculate statistical significance (ns-not significant, *P < 0.05, **P < 0.01, ***P < 0.001) between control and experimental conditions.

The 157 up-regulated genes were related to “ribose phosphate metabolic process”, “mitochondrial gene expression” and “mitochondrial translation” (**Supplementary Figure 1C**), suggesting that YAP deeply shapes and regulates mitochondrial processes, which further supports our previous data showing that mitochondrial rewiring sustains resistance to NAMPT inhibition.

The 1-phosphatidylinositol 4,5-bisphosphate phosphodiesterase beta (PLCB1; log_2_FC= - 2,46; p-value =0,006), fatty acid-binding protein (FABP4; log_2_FC=-2,34; p-value =5,27 x 10^-8^) and nicotinamide N-methyltransferase (NNMT; log_2_FC=-2,16; p-value =9,18 x 10^-5^) were amongst the most downregulated proteins in PC3 shYAP, (**Figure 3A**). Interestingly, all these proteins have been previously associated with the cellular transition towards mesenchymal phenotypes^28–33^, sustaining the role of YAP in the maintenance of stem cell-like mesenchymal phenotypes.

### NNMT is a mediator of FK866 toxicity in PC3 cells

Interestingly, NNMT is also a key enzyme in the metabolism of NAD(H)/nicotinamide (NAM)^17^. NAM is preferentially converted to nicotinamide mononucleotide (NMN) by NAMPT and consequently to NAD+, which is then regenerated to nicotinamide by NAD+-consuming enzymes as sirtuins, PARPs and other ectoenzymes as CD38. However, NAM can also be converted to 1-methyl nicotinamide (1MNA) and S-adenosylhomocysteine (SAH) by NNMT or metabolized by NAMPT to ultimately produce NAD+^34^ (**Figure 3B**).

Given the functional role of YAP and its co-downregulated protein NNMT in sustaining EMT, and the relevance of NNMT in NAD+ synthesis and signaling pathways, we investigated whether NNMT activity was also relevant for FK866 sensitivity. A strong negative correlation between viability to FK866 and NNMT expression was observed for PC cell lines (Pearson r =-0,9917; p-value = 0.0083) (**Figure 3C**). Cells silenced for NNMT (**Figure 3D-E**) were fully resistant to FK866 treatment, both in attachment-dependent and independent (soft-agar assay) conditions (**Figure 3F-G**), lacked FK866-dependent CHOP expression (**Figure 3H**), and caspase 3/7 activation (**Figure 3I**). Collectively, these data suggest that YAP-dependent NNMT induction mediates the sensitivity to NAMPT inhibition in PC3 cells.

### NNMT limits nicotinamide availability in PC3 cells

Given the fact that FK866 and NAM compete for the same binding site in the catalytic pocket of NAMPT, cellular supplementation with NAM has been previously shown to counteract FK866 toxicity by replenishing cellular NAD+ levels. In line with this, the administration of increasing concentrations of NAM to PC3 shSCR cells prevented FK866-mediated toxicity in a dose-dependent manner (**Figure 4A**).

**Figure 4.**
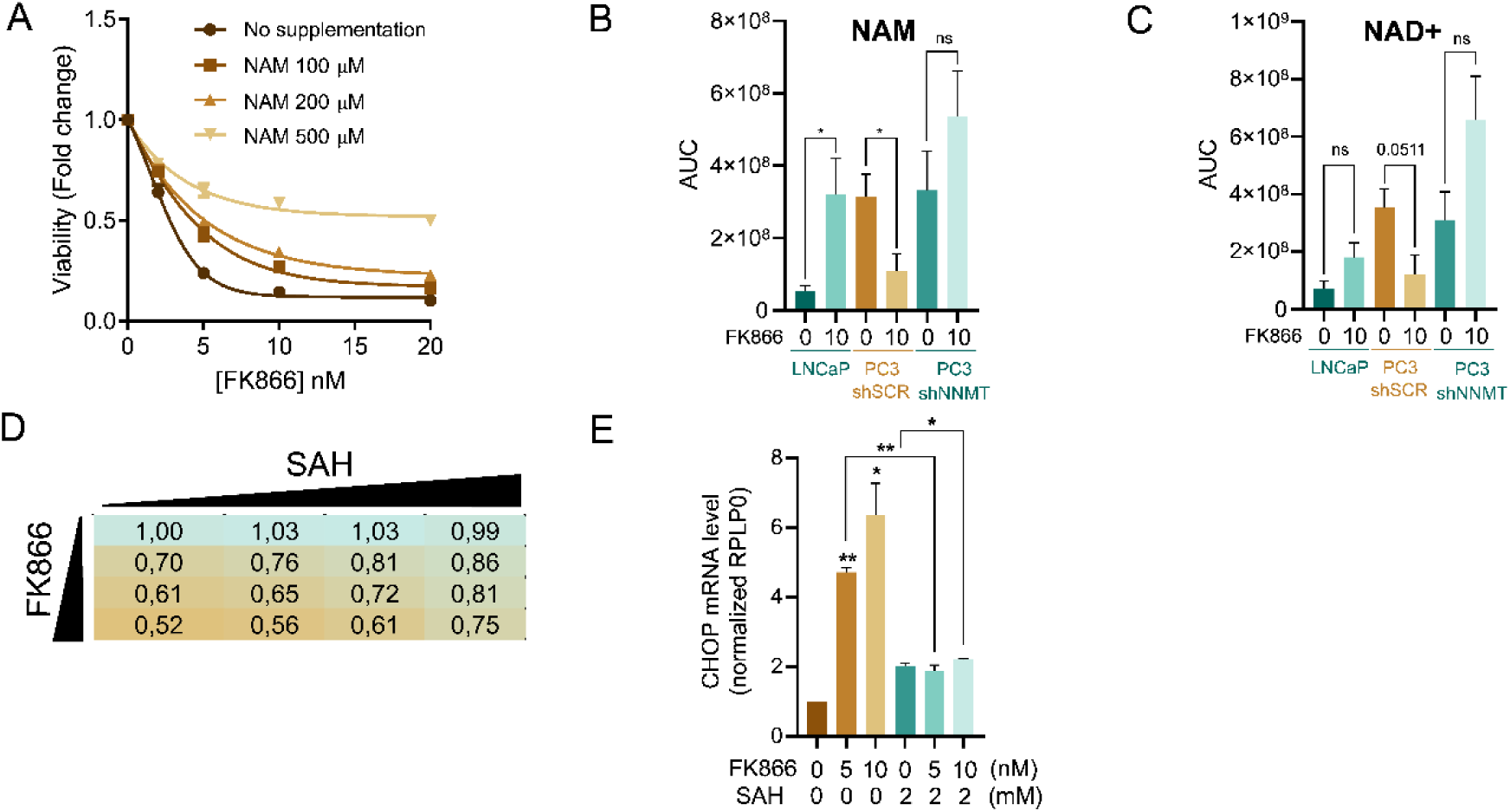
NNMT limits the availability of nicotinamide to increase FK866 toxicity**. (A)** Viability curve of PC3 cells after exposure to FK866 for 48 hours, and supplementation with NAM at the indicated concentrations (0, 100, 200 and 500 µM). Viability calculated as a fold change of the DMSO condition. Three independent biological replicates were performed**. (B)** NAM and **(C)** NAD^+^ quantification in LNCaP, PC3 shSCR and PC3 shNNMT cell lines by metabolomics analysis. Cells were exposed to 10 nM of FK866 for 48 hours, after which metabolites quantity was estimated from the area under the curve (AUC) of the spectra profile. Five independent biological replicates were performed and used for statistical analysis. **(D)** Cell viability of PC3 cells after combinatory treatment with increasing concentrations of FK866 (0, 1, 2 and 5nM) and SAH (0, 1, 2 and 5 mM). Viability is calculated as a fold change of control (DMSO). Three independent biological replicates were performed. **(E)** mRNA levels of CHOP in PC3 cells co-treated with FK866 and SAH for 48 hours, at the indicated concentrations. RPLP0 was used as housekeeping gene. Repeated measures one-way ANOVA were used to calculate statistical significance (ns-not significant, *P < 0.05, **P < 0.01, ***P < 0.001) between control and experimental conditions.

Thus, the toxic effect of NAMPT inhibition is prevented by the maintenance of the NAM cellular pool. To better understand the relevance of NNMT in preventing FK866 toxicity, we performed a mass-spectrometry analysis of the level of NAM during FK866 treatment in our cell models. Metabolomics analysis showed that PC3 shSCR treated with a non-toxic concentration of FK866 present reduced levels of NAM, likely because the available NAM is consumed by NNMT (**Figure 4B-C**). At the basal level, both PC3 shSCR and PC3 shNNMT showed a similar amount of NAM, which is explained by the higher affinity of NAM for NAMPT than NNMT. As such, the silencing of NNMT *per se* is insufficient to increase NAM levels because NAMPT-dependent NAM consumption is biochemically favored. Under NAMPT inhibition with FK866, LNCaP cells, which present very low expression of NNMT, survive FK866 toxicity by recycling the cellular NAD+ and NAM pools as NAM is not diverted to 1MNA/SAH. PC3 shNNMT cells behave similarly, showing a tendency, even though not significant, of increasing NAD+ and NAM pools upon FK866 treatment (**Figure 4B-C**). Importantly, the NNMT-derived product SAH, which inhibits NNMT activity by product accumulation, partially rescues FK866 toxicity in these cells, while fully rescuing CHOP activation upon FK866 exposure (**Figure 4D-E**).

These data suggest that decreased expression of NNMT, as observed in LNCaP and PC3 shNNMT but not in PC3 shSCR cells, can maintain the NAM pool to compete with FK866 and counteract NAMPT inhibition, and consequently sustain the cellular levels of NAD+.

### NNMT is a marker of the CRPC-SCL subtype

To further explore the clinical relevance of NNMT in prostate cancer, we investigated whether its expression is differentially regulated across clinically meaningful disease contexts. Specifically, we aimed to assess the association between NNMT expression, mesenchymal phenotype, and therapy resistance. We took advantage of the previously mentioned “CRPC-SCL” subtype dataset^24^. We analyzed the RNA-seq of the 40 samples, including prostate cancer cell lines, organoids, and patient-derived xenografts and interestingly found an upregulation of NNMT (**Figure 5A**). This observation is in line with our data, showing an upregulation of NNMT in a CRPC subtype that presents YAP activation and dependency.

**Figure 5.**
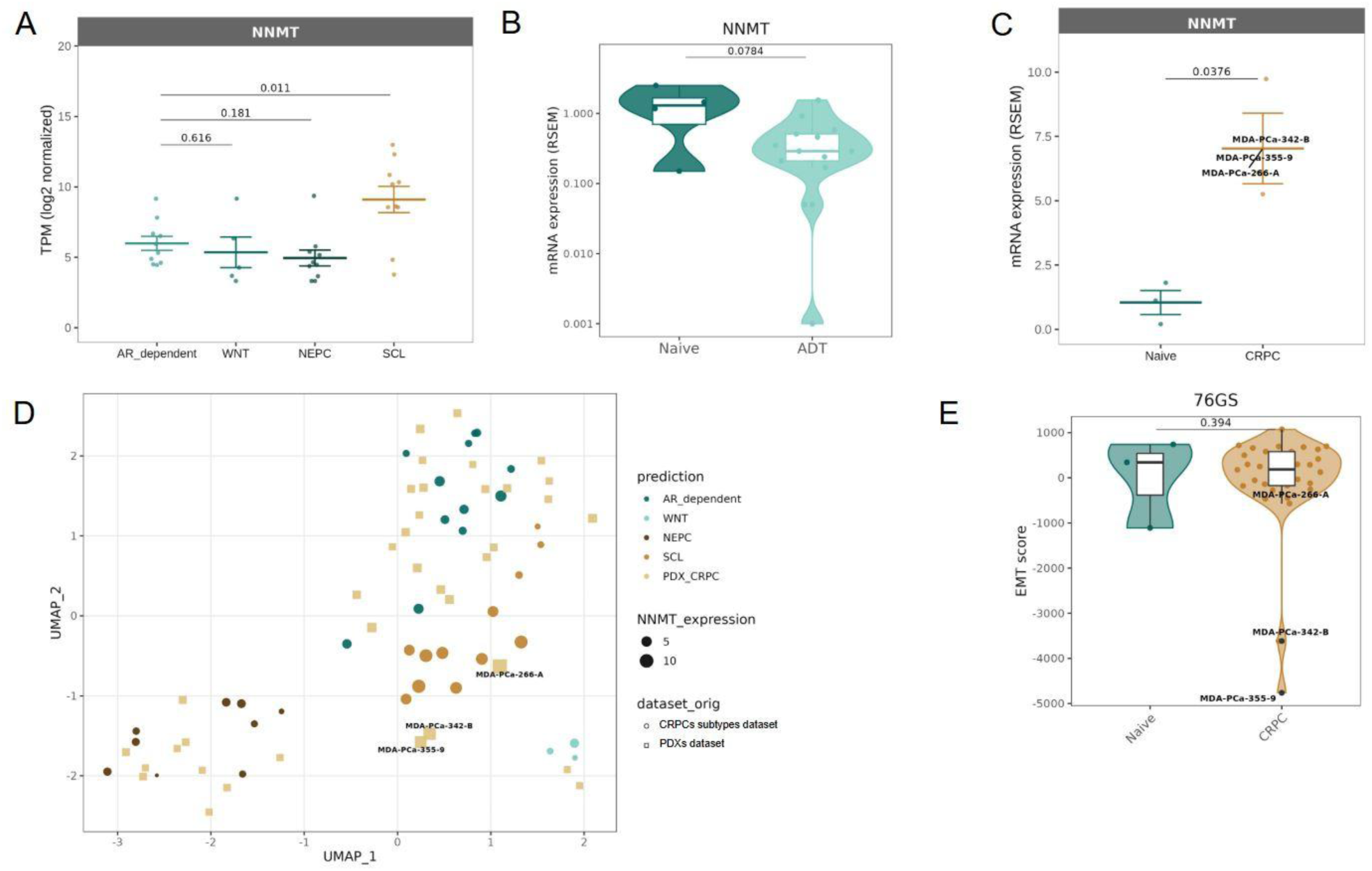
NNMT upregulation induces a stem-like phenotype in CRPC. (**A**-**C**) NNMT expression changes across CRPC subtypes and PDX samples stratified according to exposure to androgen deprivation therapy (ADT) exposure. (**D**) UMAP analysis of PDX samples integrated in CRPC subtypes dataset. Data were batch corrected after integration (**E**) Violin plots displaying EMT scores for naive and CRPC PDX samples, calculated using the 76GS scoring method. Wilcoxon rank-sum test was used to calculate statistical significance across CRPC types.

Additionally, database browsing identified a dataset of RNA-sequencing from 44 patient-derived xenograft (PDX) samples^35^. This cohort comprises models that capture a diverse range of morphological subtypes, various disease states, and multiple affected organ sites. NNMT is found to be downregulated, although not significantly, in samples deriving from patients treated with first-generation ADT (**Figure 5B**), but not in CRPC ones (**Supplementary Figure 2A**). However, if CRPC patients’ samples are classified based on the subtypes defined by ^24^, NNMT is, in fact, found upregulated in the 3 patient samples clustering with “CRPC-SCL” (**Figure 5B-C**). Moreover, it is possible to observe that also for this dataset, NNMT correlates with the different CRPC subtypes: it is highly expressed in the “CRPC-SCL” subtype and mildly expressed in all the remaining ones (**Figure 5D).** These samples presenting the highest NNMT levels are also the ones presenting the strongest EMT phenotype (**Figure 5E**).

Finally, we took advantage of data from cBioPortal (TCGA, PanCancer Atlas), and observed that patients presenting a new neoplastic event after treatment (i.e. recurrency) with leuprolide (an agent reducing testosterone levels), also showed an increased mesenchymal phenotype (**Supplementary** Figure 2B) and a concomitant significant upregulation of NNMT (Supplementary Figure 2C).

Overall, PDXs, organoids, cell lines, and patients’ data sustain a correlation between NNMT expression and the development of mesenchymal stem-cell-like phenotypes, promoting tumor aggressiveness and resistance to hormonal therapy.

### NNMT is upregulated in cancer stem-like cells with mesenchymal phenotype

Given the correlation between NNMT and CRPC-SCL, we generated a murine DVL-3 cell line that could recapitulate the stem cell-like properties (DVL3-SCM) responsible for tumor initiation, aggressiveness and therapy resistance. The DVL3 cell line had been previously isolated from prostate tumors of a Pten^-/-^/Trp53^-/-^ mouse model (DVL3-PAR)^18^. DVL3 cells were grown in low mitogen and low hormone conditions to obtain DVL3-SCM.

Transcriptomics analysis revealed a strong mesenchymal signature of DVL3-SCM compared with DVL3-PAR, as indicated by the lower calculated EMT score (**Figure 6A**). Indeed, DVL3-SCM cells presented a decreased expression of *Ck8*, *Ck18*, *Ck5* and *Ck14* cytokeratin genes, suggesting a loss of epithelial properties (**Figure 6B**). Moreover, it was observed that an increased expression of the mesenchymal gene markers *Twist*, *Slug*, *Snail* in DVL3-SCM compared with DVL3-PAR cells, along with a decrease of the epithelial marker *E-cadherin*, both at the transcript level (**Figures 6C**). Additionally, DVL3-SCM revealed an increased ability to grow in anchorage-independent (soft-agar) compared with their parental counterpart **(Figure 6D)**. The subcutaneous flank implantation of DVL3-SCM cells into wild-type C57BL/6 male mice led to increased tumor mass growth compared with DVL3-PAR implantation (**Figure 6E**), confirming DVL3-SCM increased tumorigenic potential.

**Figure 6.**
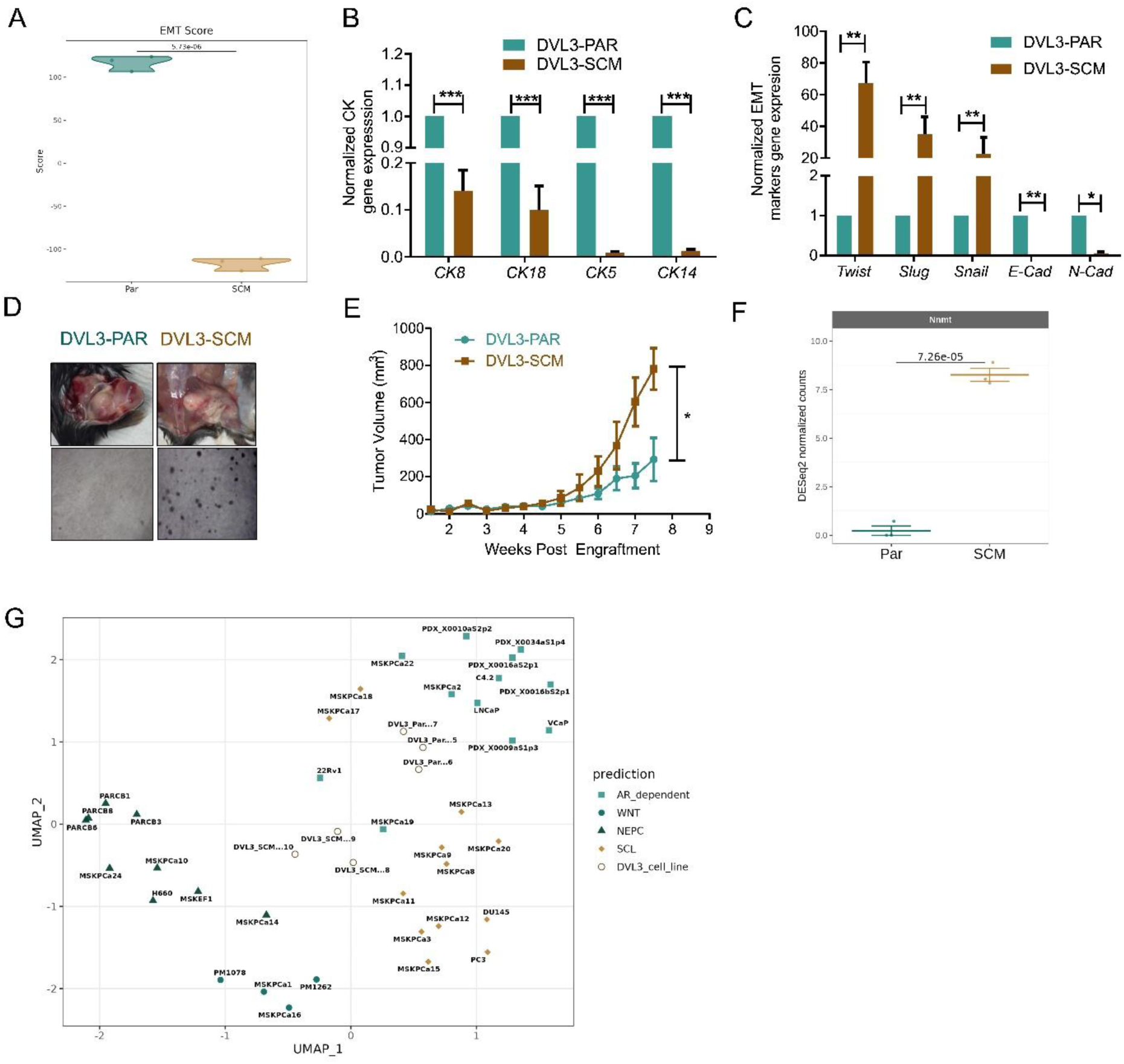
NNMT is upregulated in cancer stem-like cells with mesenchymal phenotype. (**A**) Violin plots displaying EMT scores between parental and stem cell-like murine DVL-3 cell lines, calculated using the 76GS scoring method. Expression of CK mRNAs **(B)** and EMT-associated genes **(C)** in DVL3-PAR and DVL3-SCM cells by RT-qPCR using beta-actin as the housekeeping gene. **(D)** Representative image of anchorage-independent soft-agar DVL3-PAR and DVL3-SCM cells growth. **(F)** Tumor mass volume measurements up to 8 weeks after engraftment of DVL3-PAR and DVL3-SCM cells. (**F**) NNMT expression levels between parental and stem cell-like murine samples. (**G**) UMAP visualization of the murine samples integrated with the CRPC dataset^4^. Data were batch corrected after integration. Student t-test was used to calculate statistical significance between murine samples.

Further analysis of transcriptomics data revealed NNMT (log_2_FC= 8,26; p-value =7,26 x 10^-5^) as the most upregulated transcript in DVL3-SCM cells (**Figure 6F**). Finally, we performed a UMAP analysis, integrating the RNA-seq data obtained for our murine cell lines with the one obtained by ^24^ for CRPC samples, and we observed that DVL3-SCM, differently from DVL3-PAR, cluster closer to the “CRPC-SCL” subtype, which indeed presents a NNMT upregulation (**Figure 6G**).

In conclusion, we generated a murine model that could recapitulate PC mesenchymal-stem cell-like features, including the high NNMT expression. This data confirms NNMT as a marker of the stem-cell like subtype that can be suggested for the clinical classification of CRPC patients and therapy regimens.

## DISCUSSION

Acquired resistance to NAMPT inhibition has been widely studied by us and others^8–11^. However, far less is known about the mechanisms driving and sustaining innate sensitivity to NAD(H) depleting agents. Understanding these intrinsic factors is crucial to stratify patients who may benefit from NAMPT-targeted therapies and to guide rational combination strategies. Here, we showed that YAP/NNMT axis plays a pivotal role in FK866 sensitivity in prostate cancer cells with mesenchymal, stem-like features.

The unresponsiveness of CRPC patients to the current standard of care therapies prevents successful clinical outcomes, which makes the identification of novel drug targets of critical importance. Recently, Tang and colleagues described four different classes of CRPC: AR-dependent (CRPC-AR), neuroendocrine (CRPC-NE) and the AR-null/low Wnt-dependent (CRPC-WNT) and stem cell–like (SCL) (CRPC-SCL) groups. In particular, the CRPC-SCL subtype is characterized by pronounced mesenchymal features coupled with a YAP/TAZ signature. The acquisition of mesenchymal phenotypes allows cancer cells to adapt to conditions hampering their survival, spread in the body, and resist therapy^14,15^. Interestingly, therapy-resistant PC cells presenting stem-like properties and EMT potential are particularly sensitive to NAD(H) depletion agents due to an increased expression of NAMPT, the rate-limiting enzyme of NAD^+^ synthesis pathways ^17,36^. The reason why NAMPT is upregulated in tumors presenting mesenchymal features is still unsolved, the fact that it stabilizes β-catenin^36^ is a possible explanation, nevertheless, its activity plays a critical role in cellular growth and survival under oxidative stress and chemotherapy treatments^4^.

Here, we demonstrated that the Hippo pathway effector YAP1, previously associated with EMT in diverse cancer types^37,38^, sustains the expression of proteins associated with the EMT program - PLCB1^28,29,39^, FABP4^30,31^, and NNMT-in stem cell-like CRPC models. The involvement of all three of these proteins in lipid metabolism and membrane fluidity strengthens the initial hypothesis of a crucial role of YAP in the progression of PC to the castration-resistant stage of the disease^40^ that is also achieved by fostering mesenchymal features in PC cells. Further dissection of the role of NNMT in PC tumorigenesis confirmed this enzyme as a specific marker of CRPC-SCL^24^. In keeping with that, the induction of a stem cell-like phenotype in AR-independent primary murine prostate cancer cells (DVL3-SCM) recapitulating key genetic aberrations (e.g. PTEN and TP53 double null condition) and the mesenchymal features of human PC3 cells, is enough to upregulate NNMT and induce aggressive phenotypes *in vivo*. Functionally, NNMT is closely associated with NAMPT and NAD^+^ synthesis pathways, as these enzymes compete for the substrate NAM. NAMPT and NNMT present different k_m_ for NAM binding, respectively of 1 µM and 430 µM^41^. This indicates that under physiological conditions, NAMPT binds NAM much more efficiently than NNMT, sustaining NAD^+^ synthesis. Mathematical models suggest a co-evolution of both NAMPT and NNMT, which favors NAD^+^ biosynthetic and signaling processes, as the occurrence of NNMT prevents NAM accumulation and inhibition of NAD synthesis^42^. In line with the concept that YAP-dependent control of cellular energetics can be critical in CRPC-SCL, PC cells presenting stem-like mesenchymal phenotype activate YAP transcriptional programs when exposed to the NAD(H) depleting NAMPT inhibitor FK866. Prolonged stress ultimately induces ER stress response, the transcription of CHOP, and apoptosis. Cell death is clearly dependent on NAM depletion since exogenous NAM administration, as well as NNMT silencing or its inhibition through SAH treatment, rescues FK866 lethality in CRPC-SCL cell models. Interestingly, mouse PC DVL3-SCM cells do not present an upregulation of NAMPT or a clear YAP genetic signature, leading us to the conclusion that further cellular adaptation, likely in response to the therapeutic regimen to which patients are subjected as well as metastatic processes, can also contribute to the NAMPT/NNMT oncometabolic pathway.

Notably, NAD^+^ is the donor of the ADP-ribose substrate that PARP enzymes use to modify different classes of proteins with essential roles in the regulation of cellular processes such as transcription and DNA repair. Small molecules inhibiting PARP activity have demonstrated clinical efficacy in *BRCAness* tumors and are now being tested in several types of cancer, including prostate cancer^43^. Considering that *BRCAness* cancers are often an aggressive form of tumor presenting metastatic spread already at diagnosis and poor response to treatment, the definition of NAMPT/NNMT expression and NAM metabolism may help guide clinical decision-making and contribute to maximizing the benefit of PARP inhibitor therapy through the concomitant administration of NAD^+^ depleting agents.

Overall, our data point to NNMT as a novel biomarker of the “CRPC-SCL” subtype of PC that can be used as a predictor in clinical trials aimed at evaluating the efficacy of NAD^+^-depleting agents in combination with standard therapeutic regimens.

## ETHICS STATEMENT

All procedures were performed in accordance with the Animal Scientific Procedures Act of 1986 (UK) (Project Licence Number PL2775, PL2798 and PCC943F76) which was issued by the home office. Protocols were approved by the animal welfare and ethical review body at both Queen’s University Belfast and the University of Manchester. Animals were housed in individually ventilated cages on a 12:12 light:dark cycle and *ad libitum* access to food and filtered water. Prior to each *in vivo* experiment, cells were screened for mycoplasma contamination.

## AUTHORS CONTRIBUTIONS

Conceptualization – AC, MC, AL, TT, IGM, AP; Data curation - AC, MC, NT, CMH, AdP, SLE, RS, EC, RB, DP, EF, AP; Resources – AP; Software – MC, TT; Formal analysis - AC, MC, NT, CMH, AdP, SLE, RS, EC, EF, AP; Supervision – AP; Funding acquisition – AN, AP; Validation – All; Investigation – AC, AL, AP; Visualization - AC, MC, AL, TT, IGM, AP; Methodology - AC, MC, NT, CMH, AdP, SLE, RS, DP, RB, EC, EF, MG, IC, AL, TT, IGM, AP; Writing-original draft – AC, MC, AL, AP; Project administration – AP; Writing-review and editing – All.

## SUPPLEMENTARY FIGURES

**Supplementary Figure 1:**
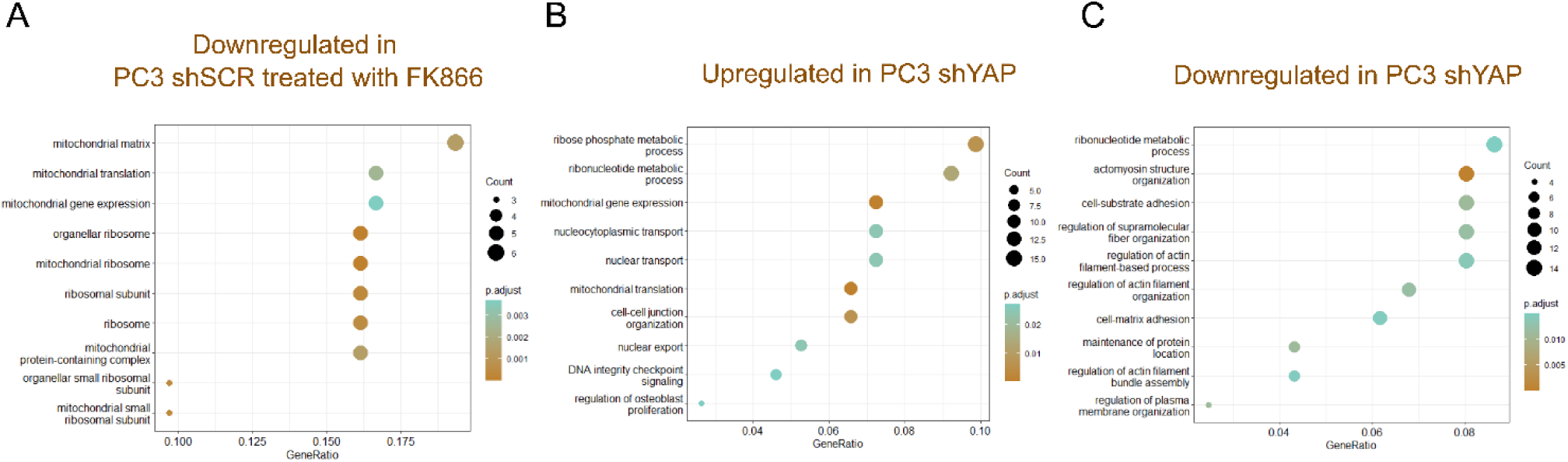
Gene set enrichment analysis (GSEA) of proteomics data. **(A)** GSEA of downregulated proteins in PC3 shSCR cell treated with FK866 (5nM) for 48 hours **(B)** GSEA of proteins up or (C) downegulated in PC3 shYAP, compared with the scramble condition (PC3 shSCR).. Significant modulated proteins presented a p-value above 0,05, calculated from three independent biological replicates.

**Supplementary Figure 2:**
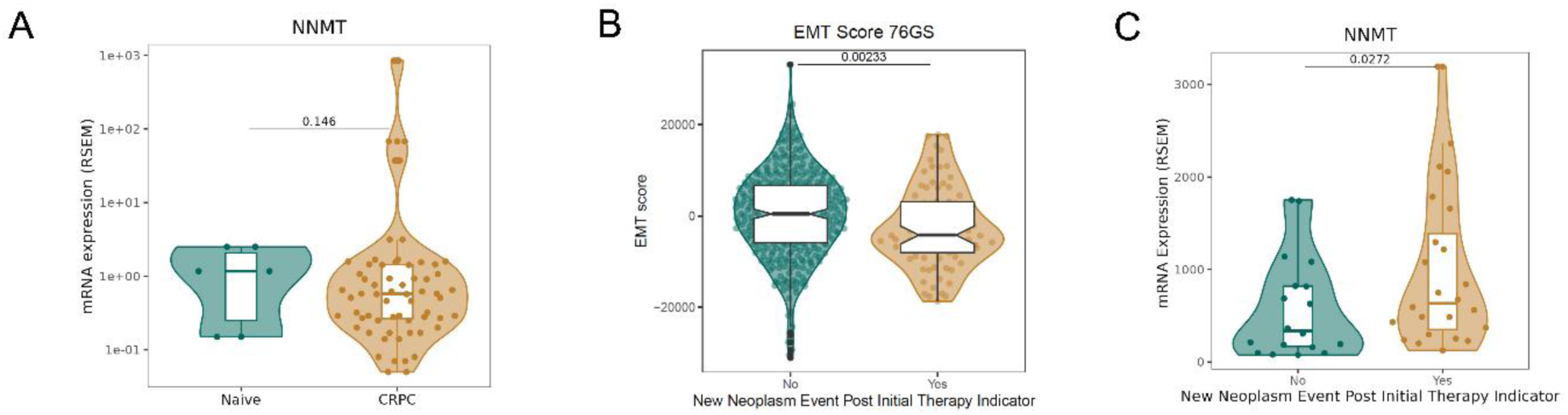
(A) NNMT expression levels of PDX samples between naive and CRPC conditions. (**B-C**) Violin plots displaying EMT scores and NNMT mRNA changes, between prostate cancer patients displaying or not a new neoplastic event after treatment. The score was calculated using the 76GS scoring method. Wilcoxon rank-sum test was used to calculate statistical significance across the different conditions.

## REFERENCES

1. Hanahan, D. Hallmarks of Cancer: New Dimensions. Cancer Discov 12, 31–46 (2022).

2. Carreira, A. S. A. et al. Mitochondrial rewiring drives metabolic adaptation to NAD(H) shortage in triple negative breast cancer cells. Neoplasia (United States) 41, 100903 (2023).

3. Zhang, P. et al. Targeting DNA damage repair functions of two histone deacetylases, HDAC8 and SIRT6, sensitizes acute myeloid leukemia to NAMPT inhibition. Clinical Cancer Research 27, 2352–2366 (2021).

4. Wang, B. et al. NAMPT overexpression in prostate cancer and its contribution to tumor cell survival and stress response. Oncogene 30, 907–921 (2011).

5. Bi, T. Q. et al. Overexpression of Nampt in gastric cancer and chemopotentiating effects of the Nampt inhibitor FK866 in combination with fluorouracil. Oncology Reports 26, 1251–1257 (2011).

6. Sawicka-Gutaj, N. et al. Nicotinamide phosphorybosiltransferase overexpression in thyroid malignancies and its correlation with tumor stage and with survivin/survivin DEx3 expression. Tumor Biology 36, 7859–7863 (2015).

7. Audrito, V. et al. Tumors carrying BRAF-mutations over-express NAMPT that is genetically amplified and possesses oncogenic properties. Journal of Translational Medicine 20, 1–13 (2022).

8. Ogino, Y., Sato, A., Uchiumi, F. & Tanuma, S. I. Cross resistance to diverse anticancer nicotinamide phosphoribosyltransferase inhibitors induced by FK866 treatment. Oncotarget 9, 16451–16461 (2018).

9. Ogino, Y. et al. Association of ABC transporter with resistance to FK866, a NAMPT inhibitor, in human colorectal cancer cells. Anticancer Res 39, 6457–6462 (2019).

10. Carreira, A. S. A. et al. Mitochondrial rewiring drives metabolic adaptation to NAD(H) shortage in triple negative breast cancer cells. Neoplasia (United States) 41, 100903 (2023).

11. Thongon, N. et al. Cancer cell metabolic plasticity allows resistance to NAMPT inhibition but invariably induces dependence on LDHA. Cancer Metab (2018) doi:10.1186/s40170-018-0174-7.

12. Bach, C. et al. The status of surgery in the management of high-risk prostate cancer. Nature Reviews Urology 11, 342–351 (2014).

13. Cheng, Q. et al. Pre-existing Castration-resistant Prostate Cancer–like Cells in Primary Prostate Cancer Promote Resistance to Hormonal Therapy. Eur Urol 81, 446–455 (2022).

14. Jiménez, N. et al. Cell Plasticity-Related Phenotypes and Taxanes Resistance in Castration-Resistant Prostate Cancer. Front Oncol 10, 1–16 (2020).

15. Zhou, J. et al. Side population rather than CD133+cells distinguishes enriched tumorigenicity in hTERT-immortalized primary prostate cancer cells. Mol Cancer 10, 1–13 (2011).

16. Tang, F. et al. Chromatin profiles classify castration-resistant prostate cancers suggesting therapeutic targets. 960, (2022).

17. Mazumder, S. et al. Integrating Pharmacogenomics Data-Driven Computational Drug Prediction with Single-Cell RNAseq to Demonstrate the Efficacy of a NAMPT Inhibitor against Aggressive, Taxane-Resistant, and Stem-like Cells in Lethal Prostate Cancer. Cancers 14, (2022).

18. Haughey, C. M. et al. Investigating radiotherapy response in a novel syngeneic model of prostate cancer. Cancers 12, 1–20 (2020).

19. Olivella, R. et al. QCloud2: An Improved Cloud-based Quality-Control System for Mass-Spectrometry-based Proteomics Laboratories. Journal of Proteome Research 20, 2010–2013 (2021).

20. Zhu, Y. et al. DEqMS: A method for accurate variance estimation in differential protein expression analysis. Molecular and Cellular Proteomics 19, 1047–1057 (2020).

21. Kim, H. J. et al. PhosR enables processing and functional analysis of phosphoproteomic data. Cell Reports 34, 108771 (2021).

22. Wu, T. et al. clusterProfiler 4.0: A universal enrichment tool for interpreting omics data. Innovation 2, 100141 (2021).

23. Perez-Riverol, Y. et al. The PRIDE database resources in 2022: A hub for mass spectrometry-based proteomics evidences. Nucleic Acids Research 50, D543–D552 (2022).

24. Tang, F. et al. Chromatin profiles classify castration-resistant prostate cancers suggesting therapeutic targets. 960, (2022).

25. Chakraborty, P., George, J. T., Tripathi, S., Levine, H. & Jolly, M. K. Comparative Study of Transcriptomics-Based Scoring Metrics for the Epithelial-Hybrid-Mesenchymal Spectrum. Frontiers in Bioengineering and Biotechnology 8, 1–13 (2020).

26. Eddie, S. L. et al. Tumorigenesis and peritoneal colonization from fallopian tube epithelium. Oncotarget 6, 20500–20512 (2015).

27. Adams, K. J. et al. Skyline for Small Molecules: A Unifying Software Package for Quantitative Metabolomics. Journal of Proteome Research 19, 1447–1458 (2020).

28. Liang, S. et al. A PLCB1-PI3K-AKT Signaling Axis Activates EMT to Promote Cholangiocarcinoma Progression. Cancer Research 81, 5889–5903 (2021).

29. Wang, Y. et al. PLCB1 Enhances Cell Migration and Invasion in Gastric Cancer Via Regulating Actin Cytoskeletal Remodeling and Epithelial–Mesenchymal Transition. Biochemical Genetics 61, 2618–2632 (2023).

30. Furuhashi, M., Saitoh, S., Shimamoto, K. & Miura, T. Fatty acid-binding protein 4 (FABP4): Pathophysiological insights and potent clinical biomarker of metabolic and cardiovascular diseases. Clinical Medicine Insights: Cardiology 2014, 23–33 (2014).

31. Tian, W. et al. FABP4 promotes invasion and metastasis of colon cancer by regulating fatty acid transport. Cancer Cell International 20, 1–13 (2020).

32. Wang, Y. et al. NNMT contributes to high metastasis of triple negative breast cancer by enhancing PP2A/MEK/ERK/c-Jun/ABCA1 pathway mediated membrane fluidity. Cancer Letters 547, 215884 (2022).

33. Hah, Y. S. et al. Nicotinamide N-methyltransferase induces the proliferation and invasion of squamous cell carcinoma cells. Oncology Reports 42, 1805–1814 (2019).

34. Hah, Y. S. et al. Nicotinamide N-methyltransferase induces the proliferation and invasion of squamous cell carcinoma cells. Oncol Rep 42, 1805–1814 (2019).

35. Anselmino, N. et al. Integrative Molecular Analyses of the MD Anderson Prostate Cancer Patient-derived Xenograft (MDA PCa PDX) Series. Clinical Cancer Research 30, 2272–2285 (2024).

36. Lee, J. et al. Selective Cytotoxicity of the NAMPT Inhibitor FK866 Toward Gastric Cancer Cells With Markers of the Epithelial-Mesenchymal Transition, Due to Loss of NAPRT. Gastroenterology 155, 799–814.e13 (2018).

37. Cheng, D., Jin, L., Chen, Y., Xi, X. & Guo, Y. YAP promotes epithelial mesenchymal transition by upregulating Slug expression in human colorectal cancer cells. International journal of clinical and experimental pathology 13, 701–710 (2020).

38. Yuan, Y. et al. YAP overexpression promotes the epithelial-mesenchymal transition and chemoresistance in pancreatic cancer cells. Molecular Medicine Reports 13, 237–242 (2016).

39. Sun, L. et al. ENO1 promotes liver carcinogenesis through YAP1-dependent arachidonic acid metabolism. Nature Chemical Biology 19, 1492–1503 (2023).

40. Lee, H. C. et al. YAP1 overexpression contributes to the development of enzalutamide resistance by induction of cancer stemness and lipid metabolism in prostate cancer. Oncogene 40, 2407– 2421 (2021).

41. Wang, W., Yang, C., Wang, T. & Deng, H. Complex roles of nicotinamide N-methyltransferase in cancer progression. Cell Death and Disease 13, (2022).

42. Bockwoldt, M. et al. Identification of evolutionary and kinetic drivers of NAD-dependent signaling. Proceedings of the National Academy of Sciences of the United States of America 116, 15957– 15966 (2019).

43. Maria Teresa Bourlon, P. V. and E. C. Development of PARP inhibitors in advanced prostate cancer. Therapeutic Advances in Medical Oncology 9, 259–261 (2024).

